# Early Life Supraphysiological Levels of Oxygen Exposure Permanently Impairs Hippocampal Mitochondrial Function

**DOI:** 10.1101/668111

**Authors:** Manimaran Ramani, Kiara Miller, Ranjit Kumar, Jegen Kadasamy, Lori McMahon, Scott Ballinger, Namasivayam Ambalavanan

## Abstract

Preterm infants requiring prolonged oxygen therapy often develop cognitive dysfunction later life. Previously, we reported that 14-week-old young adult mice exposed to hyperoxia as newborns had spatial memory deficits and hippocampal shrinkage. We hypothesized that the underlying mechanism was the induction of hippocampal mitochondrial dysfunction by neonatal hyperoxia. C57BL/6j mouse pups were exposed to 85% oxygen or air from P2 - P14. Hippocampal proteomic analysis was performed in young adult mice (14 weeks). Mitochondrial bioenergetics were measured in neonatal (P14) and young adult mice. We found that hyperoxia exposure reduced mitochondrial ATP-linked oxygen consumption and increased state 4 respiration linked proton leak in both neonatal and young adult mice. Following hyperoxia exposure, complex I function was decreased at P14 but increased in young adult mice. Proteomic analysis revealed that neonatal hyperoxia exposure decreased complex I NDUFB8 and NDUFB11 and complex IV 7B subunits, but increased complex III subunit 9 in young adult mice. In conclusion, neonatal hyperoxia permanently impairs hippocampal mitochondrial function and alters complex I function. These changes may account for memory deficits seen in preterm survivors following prolonged oxygen supplementation and may potentially be a contributing mechanism in other oxidative stress associated cognitive disorders.

## Introduction

Many extremely preterm infants often require prolonged periods of supraphysiological oxygen (hyperoxia) exposure due to lung immaturity. In addition, even preterm infants not receiving supplemental oxygen are exposed to a relatively hyperoxemic environment compared to the hypoxemic normal intrauterine environment (PO_2_ 25-35 mm Hg) during a critical developmental period for many organ systems. Preterm infants requiring prolonged oxygen supplementation are at higher risk of morbidities such as retinopathy of prematurity^1,2^ and chronic lung disease (bronchopulmonary dysplasia [BPD])^3,4^, probably as a consequence of chronic oxidative stress (OS). Children with BPD frequently exhibit deficits in executive function and cognition even in the absence of apparent brain injuries such as intraventricular hemorrhage and periventricular leukomalacia^5–8^. Although direct effects of OS and lung injury-induced systemic inflammation on the developing brain have been considered as possible etiologies, the exact mechanism(s) by which children with BPD develop cognitive dysfunction despite no apparent brain injury is not known.

Although the long-term detrimental effects of early hyperoxia exposure on lung development and function have been studied extensively, little is known about its effect on brain development and function. Previously, we have shown that in C57BL/6J mice, hyperoxia (85% oxygen [O_2_]) exposure during the neonatal period (P2-14) (neonatal hyperoxia) leads to spatial memory and learning deficits, increased exploratory behavior, and shrinkage of area CA1 of the hippocampus when assessed at young adult age (14 weeks)^9^. Recently, our proteomic analysis of hippocampal homogenates from neonatal mice (P14) exposed to hyperoxia from P2-14 indicated impairments in hippocampal protein synthesis and translation and predicited mitochondrial dysfunction^10^. Hyperoxic exposure can cause cell death^11^ and impair cell survival^12^ in the developing brain. Chronic OS following O_2_ supplementation may negatively affect neuronal mitochondrial function and lead to neurodegenerative disorders^13^. Area CA1 of the hippocampus, a region of the hippocampus essential for the acquisition of long-term memory^14–16^, is highly vulnerable to OS^17^. Mitochondria isolated from hippocampal CA1 neurons release more reactive oxygen species than other regions of the hippocampus^18^. Adequate mitochondrial function is essential for mechanisms required for learning and memory^19^ such as maintenance of long-term synaptic plasticity (strengthening of synapses; long-term potentiation, and weakening of synapses; long-term depression) in the hippocampus. While mitochondrial dysfunction is associated with the pathogenesis of several neurodegenerative diseases in adults^20^, the impact of early-life mitochondrial dysfunction on long-term brain development and function is not known.

In this study, we hypothesized that prolonged hyperoxia exposure during the critical developmental period would permanently alter hippocampal mitochondrial function. Our objective was to determine the long-term changes in hippocampal mitochondrial respiratory complex protein expression and bioenergetic function in neonatal mice (P14) and young adult mice (14 weeks) exposed to hyperoxia from P2-P14.

## Results

### Targeted and Global Proteomics

#### Long-term Effect of Neonatal Hyperoxia Exposure on Hippocampal Mitochondrial Complex I, II, and III Protein Expressions in Young Adult Mice (14 weeks)

Young adult mice exposed to hyperoxia as neonates had reduced amounts of complex I NADH Dehydrogenase [Ubiquinone] 1 Beta Subcomplex 8 (NDUFB8), and complex I NADH Dehydrogenase [Ubiquinone] 1 Beta Subcomplex 11 (NDUFB11) subunit proteins (Table 1). The levels of other detected complex I subunits and of complex II subunits were comparable between the groups. (Table 1). Complex III cytochrome b-c1 complex subunit 9 was increased in the hyperoxia-exposed group compared to air-exposed group (Table 1).

**Table 1:** Long-term Effect of Neonatal Hyperoxia Exposure on Hippocampal Mitochondrial Complex I, II and III Proteins in Young Adult Mice (n=5 in Air group, 6 in Hyperoxia group)

| Molecule (Symbol) | Protein Log Fold Change in Hyperoxia (vs. Air) | P value for protein change |
| --- | --- | --- |
| **Complex I Protein Subunits** |  |  |
| NADH Dehydrogenase [Ubiquinone] 1 Alpha Subcomplex |  |  |
| Subunit 2 (NDUFA2) | -0.27 | 0.81 |
| Subunit 5 (NDUFA5) | +0.65 | 0.38 |
| Subunit 6 (NDUFA6) | -0.18 | 0.90 |
| Subunit 7 (NDUFA7) | +1.34 | 0.23 |
| Subunit 8 (NDUFA8) | +2.19 | 0.08 |
| Subunit 9 (NDUFA9) | +0.82 | 0.51 |
| Subunit 10 (NDUFA10) | -0.54 | 0.65 |
| Subunit 12 (NDUFA12) | +1.36 | 0.36 |
| Subunit 13 (NDUFA13) | +1.29 | 0.26 |
| Assembly Factor 3 (NDUFAF3) | -0.90 | 0.58 |
| Assembly Factor 4 (NDUFAF4) | +0.26 | 0.85 |
| Assembly Factor 5 (NDUFAF5) | +0.09 | 0.94 |
| NADH Dehydrogenase [Ubiquinone] 1 Beta Subcomplex |  |  |
| Subunit 3 (NDUFB3) | +0.02 | 0.98 |
| Subunit 4 (NDUFB4) | -0.35 | 0.79 |
| Subunit 5 (NDUFB5) | +0.42 | 0.78 |
| Subunit 7 (NDUFB7) | +1.10 | 0.46 |
| **Subunit 8 (NDUFB8)** | **-2.65** | **0.04** |
| Subunit 10 (NDUFB10) | +0.37 | 0.63 |
| **Subunit 11, (NDUFB11)** | **-2.37** | **0.034** |
| NADH Dehydrogenase [Ubiquinone] 1 |  |  |
| Subunit C2 (NDUFC2) | +0.07 | 0.95 |
| NADH Dehydrogenase [Ubiquinone] |  |  |
| Flavoprotein 1 (NDUFV1) | +1.94 | 0.17 |
| Flavoprotein 2 (NDUFV2) | +0.54 | 0.19 |
| **Complex II Protein Subunits** |  |  |
| Succinate dehydrogenase |  |  |
| Cytochrome b560 subunit (SDHC) | +0.89 | 0.53 |
| Iron-sulfur subunit (SDHB) | +0.82 | 0.49 |
| Flavoprotein subunit (SDHA) | -0.14 | 0.64 |
| **Complex III Protein Subunits** |  |  |
| Cytochrome B5 Type B (CYB5B) | -0.38 | 0.78 |
| Cytochrome b-c1 complex subunit 9 (UQCR10) | **+4.44** | **0.0005** |

#### Long-term Effect of Neonatal Hyperoxia Exposure on Hippocampal Mitochondrial Complex IV and V Protein Expressions in Young Adult Mice

Young adult mice exposed to hyperoxia as neonates had less cytochrome C oxidase subunit 7B (COX7B) protein compared to air-exposed groups (Table 2). The amounts of other detected complex IV and V subunits were comparable between the groups (Table 2).

**Table 2:** Long-term Effect of Neonatal Hyperoxia Exposure on Hippocampal Mitochondrial Complex IV and V Proteins in Young Adult Mice (n=5 in Air group, 6 in Hyperoxia group)

| Molecule (Symbol) | Protein Log Fold Change in Hyperoxia (vs. Air) | P Value for Protein Change |
| --- | --- | --- |
| **Complex IV Protein Subunits** |  |  |
| Cytochrome C Oxidase |  |  |
| Subunit 2 (COX2) | +0.67 | 0.06 |
| Subunit 4 Isoform 1 (COX4I) | +0.42 | 0.59 |
| Subunit 5A (COX5A) | -1.42 | 0.33 |
| Subunit 5B (COX5B) | +0.15 | 0.78 |
| Subunit 6A1 (CX6A1) | +3.32 | 0.08 |
| Subunit 6B1 (CX6B1) | +0.55 | 0.34 |
| Subunit 6C (COX6C) | +0.19 | 0.87 |
| Subunit 7A2 (CX7A2) | +0.96 | 0.51 |
| **Subunit 7B (COX7B)** | **-2.09** | **0.04** |
| Subunit 7C (COX7C) | +0.39 | 0.39 |
| Assembly Factor 7 (COA7) | +1.76 | 0.18 |
| Subunit NDUFA4 (NDUA4) | +1.17 | 0.06 |
| Translational Activator 1 (TACO1) | -0.74 | 0.58 |
| **Complex V Protein Subunits** |  |  |
| ATP Synthase |  |  |
| Protein 8 (ATP8) | +0.66 | 0.25 |
| Subunit Alpha (ATPA) | +0.16 | 0.34 |
| Subunit Beta (ATPB) | +0.02 | 0.93 |
| Subunit Delta (ATPD) | -0.34 | 0.78 |
| Subunit Gamma (ATPG) | +0.44 | 0.48 |
| Subunit F (ATPK) | -1.82 | 0.30 |
| Subunit O (ATPO) | +0.47 | 0.23 |
| Subunit D (ATP5H) | +0.73 | 0.06 |
| Subunit G (ATP5L) | -0.33 | 0.85 |
| Subunit E (ATP5I) | +0.89 | 0.11 |
| Subunit Epsilon (ATP5E) | -1.35 | 0.36 |
| Subunit S (ATP5S) | +0.88 | 0.37 |

### Bioinformatic Analysis of Differentially Expressed Hippocampal Proteins

#### Differentially Expressed Hippocampal Proteins in Young Adult Mice Following Neonatal Hyperoxia Exposure

Using a cut-off of ±1.5 fold-change with P-value <0.05 (by analysis of variance), and a false discovery rate of 5%, we identified 196 hippocampal proteins that were differentially expressed in the neonatal hyperoxia-exposed young adult mice compared to air-exposed young adult mice. Of these 196 proteins, 48 proteins were increased, and 148 were decreased following neonatal hyperoxia exposure. The heat map of the differentially expressed proteins is shown in Supplemental Fig 1. The full list of upregulated and downregulated proteins (limited to fold-change 1.5) in young adult mice exposed to neonatal hyperoxia are listed in **Supplemental Tables 1** and **2**, respectively. The top 10 differentially expressed proteins in the hyperoxia-exposed group are listed in Table 3. The protein classes that are upregulated and downregulated by hyperoxia exposure are shown in Fig 1A and 1B, respectively.

**Table 3:**
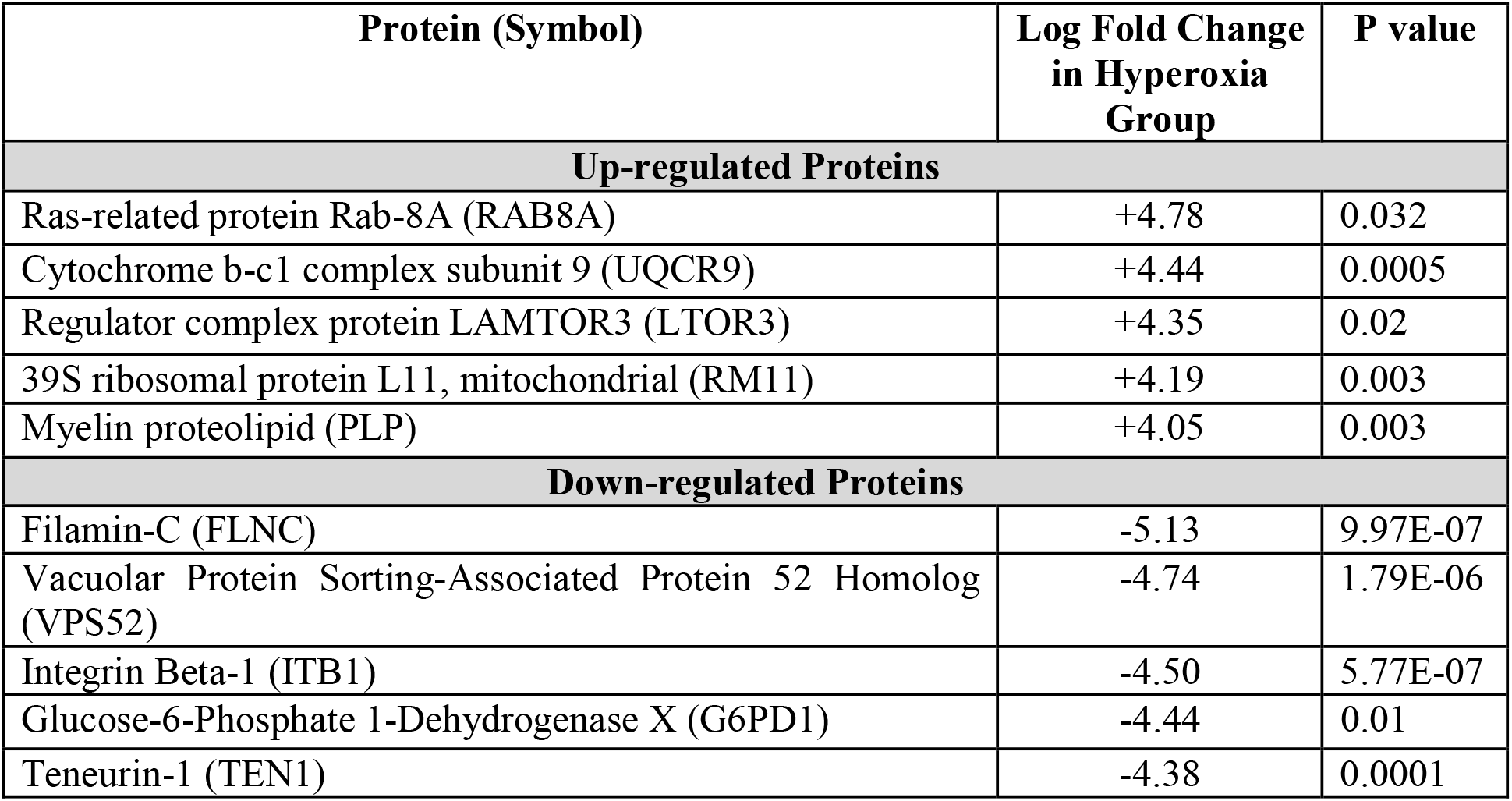
Top 10 Differentially Expressed Hippocampal Proteins in Neonatal Hyperoxia-Exposed Young Adult Mice (n=5 in Air group, 6 in Hyperoxia group, P= Hyperoxia vs. Air group)

**Fig 1:**
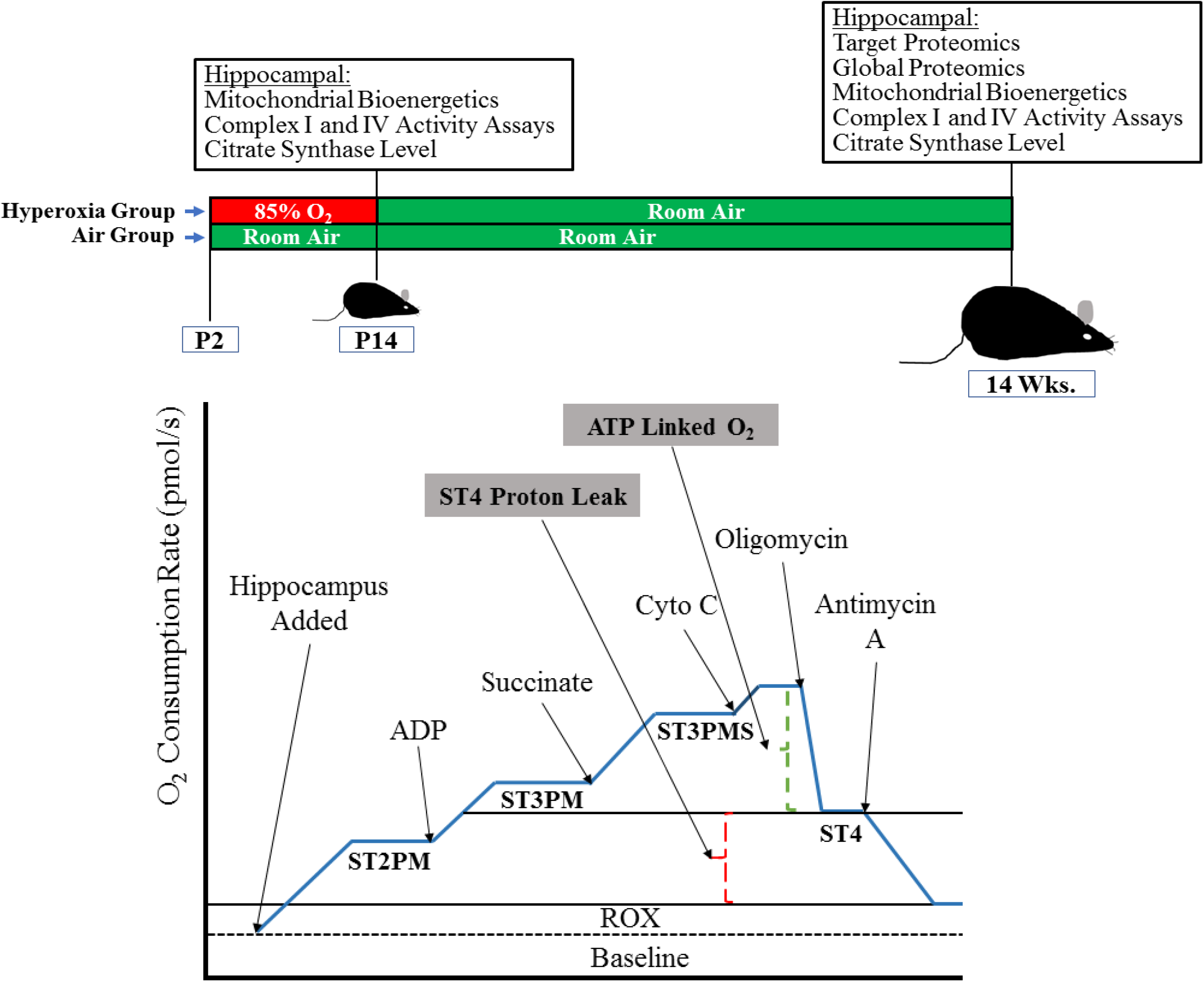
Distribution of hippocampal proteins in young adult mice exposed to neonatal hyperoxia. A. Graphical representation of the distribution of upregulated hippocampal proteins by class in the hyperoxia-exposed group. B. Graphical representation of the distribution of downregulated hippocampal proteins by class in the hyperoxia-exposed group. C. Graphical representation of the distribution of upregulated hippocampal proteins by biological processes in the hyperoxia-exposed group. D. Graphical representation of the distribution of downregulated hippocampal proteins by biological processes in the hyperoxia-exposed group.

#### Long-term Effect of Effect of Neonatal Hyperoxia Exposure on Hippocampal Biological Processes in Young Adult Mice

Differentially expressed hippocampal proteins were predominantly involved in biological processes of cellular process, metabolic process, biogenesis, and protein localization (Fig 1C and 1D). Among the upregulated proteins, functions of 20 (29.9%) proteins were related to cellular process, 18 (26.9%) were proteins related to metabolic process, 9 (13.4%) were proteins related to biogenesis process, and 6 (9%) were proteins related to localization process (Fig 1C). Among the downregulated proteins, functions of 62 (27.8%) proteins were related to cellular process, 39 (18.1%) were proteins related to metabolic process, 26 (12.1%) were proteins related to biogenesis process, and 20 (9.6%) were proteins related to localization process (Fig 1D).

#### Top Canonical Hippocampal Pathways Regulated by Neonatal Hyperoxia Exposure in Young Adult Mice

The top canonical pathways that were impacted by neonatal hyperoxia exposure are listed in Table 4. Bioinformatic analysis of differentially expressed proteins predicated that mitochondrial function (P = 2.03E-06), oxidative phosphorylation (P = 1.25E-05), GABA receptor signaling (P = 1.26E-04), amyotrophic lateral sclerosis signaling (P = 4.09E-04), and amyloid processing (P = 5.34E-04) were impacted in young adult mice that had neonatal hyperoxia exposure. Fifteen proteins were related to mitochondrial function, 11 were related to oxidative phosphorylation, 9 were related to GABA receptor signaling, 9 were proteins related to amyotrophic lateral sclerosis signaling, and 6 were proteins related to amyloid processing and were differentially expressed in the hyperoxia-exposed group.

**Table 4:** Top Canonical Pathways Involved in Neonatal Hyperoxia-Exposed Young Adult Mice by Ingenuity Pathway Analysis (Using Proteomics Data, n=5 in Air group, 6 in Hyperoxia group, P= Hyperoxia vs. Air group)

| Name | P-Value | Overlap<br>(percentage; number of proteins<br>differentially expressed/number<br>of proteins in pathway) |
| --- | --- | --- |
| Mitochondrial dysfunction | 2.03E-06 | 8.8; 15/171 |
| Oxidative phosphorylation | 1.25E-05 | 10.1; 11/109 |
| GABA receptor signaling | 1.26E-04 | 9.5; 9/95 |
| Amyotrophic lateral sclerosis signaling | 4.09E-04 | 8.1; 9/111 |
| Amyloid processing | 5.34E-04 | 11.8; 6/51 |

### Mitochondrial Studies

#### Effects of Neonatal Hyperoxia Exposure on Hippocampal Mitochondrial Bioenergetics in Neonatal Mice (P14)

Neonatal hyperoxia (exposure from P2-P14) exposure decreased both pyruvate/malate mitochondrial ATP linked (P = 0.01) and complex I enzyme activity (P = 0.01) at P14 (Fig 2A and 2D, respectively). No differences were observed between hyperoxia-exposed and air-exposed controls in ATP linked O_2_ consumption rates that utilized succinate (complex II substrate; P = 0.73) or complex IV activity (P = 0.99) (Fig 2C and 2E, respectively), consistent with the hypothesis that the observed differences were related to a complex I defect. Examination of state 4 minus basal O_2_ consumption rates also suggested increased oxygen consumption in the hyperoxia-exposed group (P = 0.03), which could be linked to increased proton leak and/or oxidant generation (Fig 2B). Hyperoxia-exposed neonatal mice also had reduced citrate synthase activity (P = 0.01) (Fig 2F), compared to air-exposed neonatal mice.

**Fig 2:**
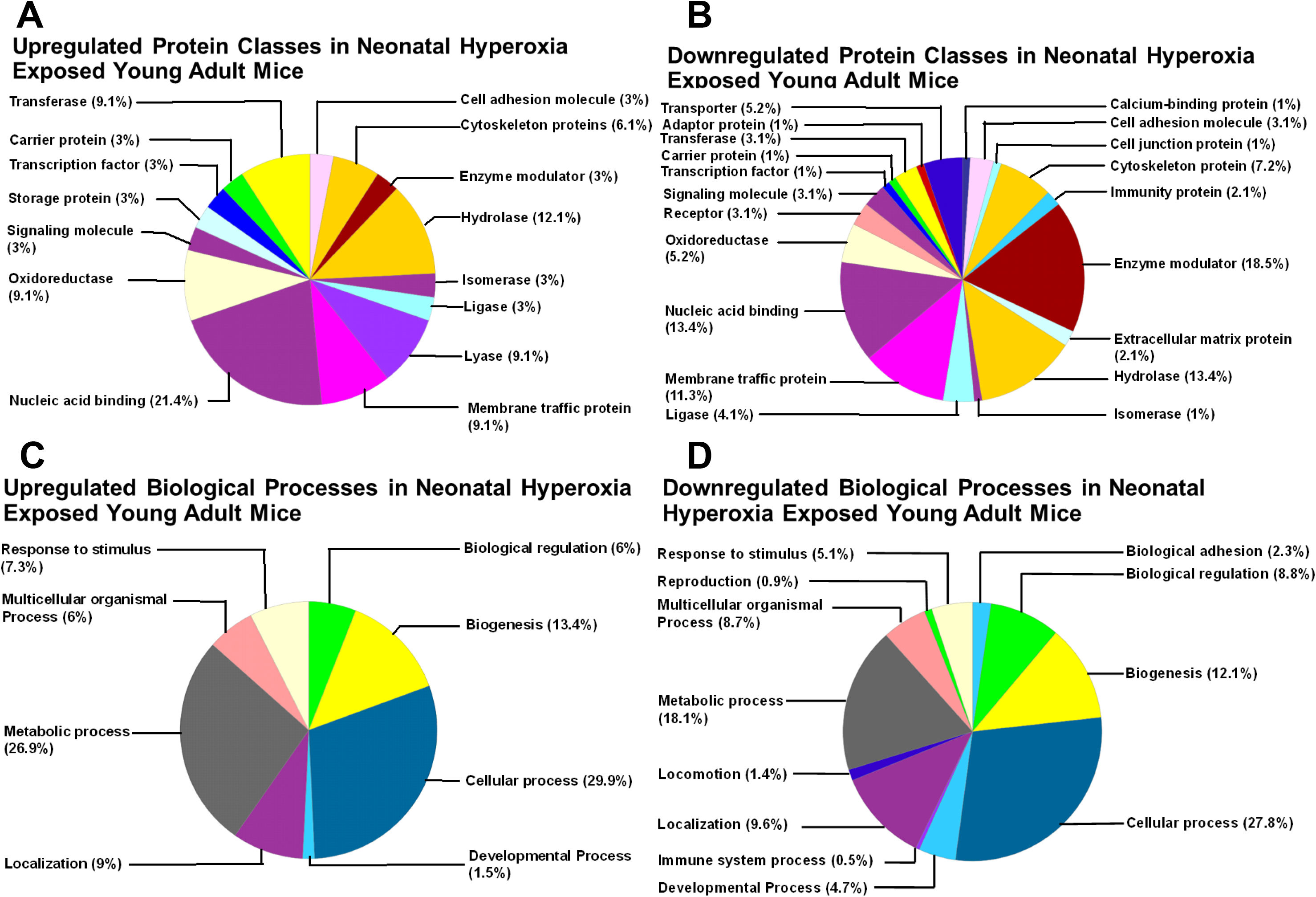
Effects of neonatal hyperoxia exposure on hippocampal mitochondrial bioenergetics in neonatal mice. A = ATP Linked Oxygen Consumption, B = State 4 Respiration Proton Leak, C = Succinate Induced Oxygen Consumption, D = Complex I Activity measured by assay, E = Complex IV Activity measured by assay, and F = Citrate Synthase Activity measure by assay,. Air-exposed: cyan bars with horizontal stripes and hyperoxia-exposed: solid red bars; means± SEM; n= 9 in air and 9 in hyperoxia.*p<0.05 vs. air-exposed mice.

#### Effects of Neonatal Hyperoxia Exposure on Hippocampal Mitochondrial Bioenergetics in Young Adult Mice (14 weeks)

Similar to the observations in neonates exposed to hyperoxia, adult mice (14 weeks old) that underwent neonatal hyperoxia exposure from P2-P14 had decreased mitochondrial ATP linked O_2_ consumption in the presence of complex I substrates (pyruvate/malate) (P = 0.01) (Fig 3A) whereas in the presence of succinate (P = 0.74), no differences were observed relative to air-exposed controls (Fig 3D). Similarly, no differences were observed in complex IV activity (P= 0.45) between exposed and unexposed control groups (Fig 3G). Also, similar to the hyperoxia-exposed newborn mice, oligomycin induced state 4 O_2_ consumption rates minus basal O_2_ consumption rates were increased (P = 0.05) (Fig 3C), consistent with increased proton leak and/or oxidant generation. However, unlike in neonates, complex I activity (P = 0.002) was significantly increased in the 14-week old mice exposed to hyperoxia as neonates (Fig 3E). Subgroup analysis by sex also showed that young adult male mice exposed to hyperoxia as neonates had decreased ATP linked O_2_ consumption (One Way ANOVA, mean difference 36.7, P = 0.01) (Fig 3B) and increased complex I activity (One Way ANOVA, mean difference 13.66, P = 0.03) (Fig 3F) compared to air-exposed young adult male mice. The difference in citrate synthase activity (P = 0.78) seen in neonatal mice exposed to hyperoxia were no longer observed when assessed as young adults (Fig 3H).

**Fig 3:**
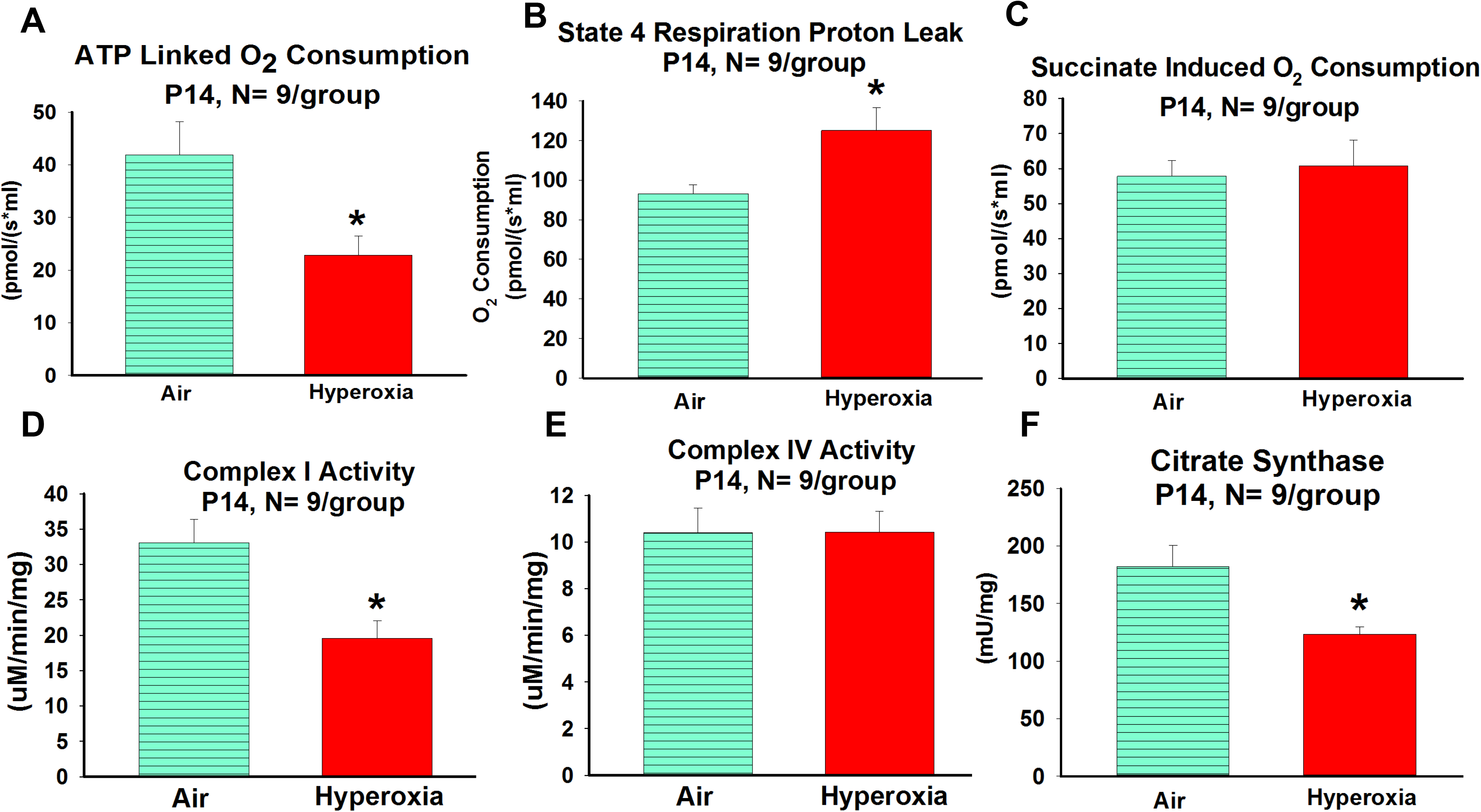
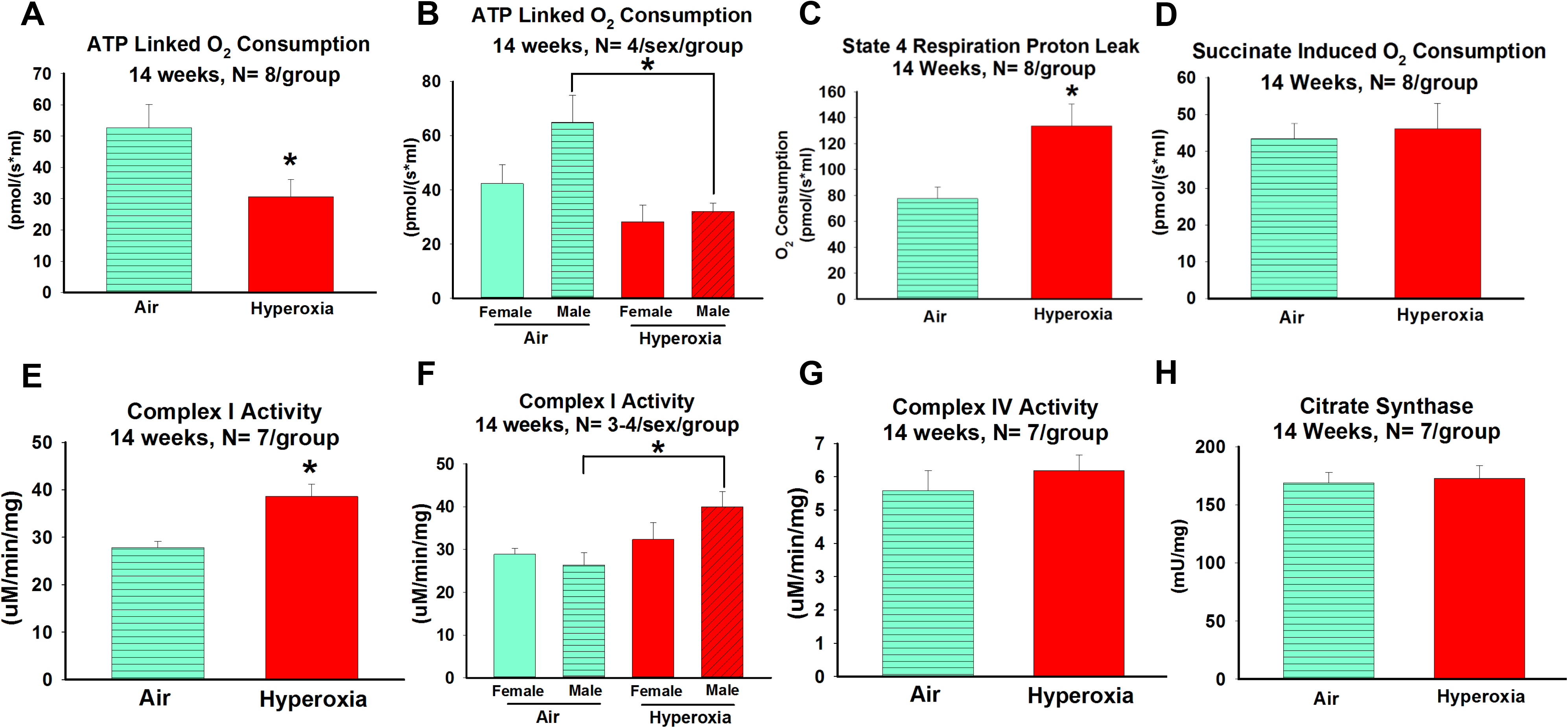
Effects of neonatal hyperoxia exposure on hippocampal mitochondrial bioenergetics in young adult mice. A = ATP Linked Oxygen Consumption, B = ATP Linked Oxygen Consumption by Sex, C = State 4 Respiration Proton Leak, D = Succinate Induced Oxygen Consumption, E = Complex I Activity measured by assay, F = Complex I Activity by Sex, G = Complex IV Activity measured by assay, and H = Citrate Synthase Activity measure by assay; no difference between the groups. Fig A, C, D, E, G, and G; air-exposed: cyan bars with horizontal stripes and hyperoxia-exposed: solid red bars; means± SEM; n= 7-8 in air and 7-9 in hyperoxia.*p<0.05 vs. air-exposed mice. Fig B, and F; air-exposed females: solid cyan bars, air-exposed males: cyan bars with horizontal stripes, hyperoxia-exposed females: solid red bar, and hyperoxia-exposed males, red bars with angled stripes; means± SEM; n= 4/sex/group.*p<0.05 = air-exposed mice vs. hyperoxia-exposed males by One Way ANOVA.

## Discussion

This is the first preclinical study to demonstrate the long-term adverse effect of early life hyperoxia on hippocampal mitochondrial function and mitochondrial respiratory chain protein expression. We discovered that hyperoxia exposure during a critical developmental period permanently impairs hippocampal mitochondrial function, alters the expression of specific respiratory chain subunits for complexes I and III, and impairs complex I activity in the hippocampus. As spatial memory deficits and other cognitive problems in the mouse model of bronchopulmonary dysplasia (BPD) correspond to the cognitive deficits seen in adolescents with BPD, these new observations suggest that permanent hippocampal mitochondrial dysfunction induced by early life oxygen exposure as a contributor to the pathophysiology of BPD associated cognitive dysfunction.

Our study has the strength of unbiased proteomic analysis of whole hippocampal tissue using highly sensitive mass spectrometric methods. Rather than being limited to only mitochondrial proteins, our study also evaluated the long-term impact of early life oxygen exposure on all hippocampal proteins and used sophisticated bioinformatics analysis to define long-term changes in the hippocampal signaling pathways. This study also evaluated mitochondrial bioenergetics induced by hyperoxia exposure during the critical developmental period (P14) and young adult age (14 weeks, the age at which we observed cognitive dysfunction in our previous study).

There are certain limitations to this study. The proteomics and the mitochondrial bioenergetic studies were performed on the whole hippocampus instead of specific hippocampal subfields which are known to play different roles in learning and memory. Furthermore, since proteomic and bioenergetic studies were done from the whole hippocampus, it is not possible to determine whether these oxygen-induced proteomic and mitochondrial functional changes were predominantly derived from neurons or glial cells or a combination of both. In this study, we focused on proteomics and mitochondrial bioenergetics only from the hippocampal homogenates, and not from regions such as cerebellum, amygdala, corpus callosum, and white matter tracts which may also are impacted by hyperoxia exposure^21–23^. Even though hippocampal complexes I and IV activities were measured, complex III and complex V activities were not measured due to technical difficulties and the size of the hippocampus.

Though lung and brain development in newborn mouse pups corresponds to 24-28 weeks of gestation in human preterm infants, the highly efficient redox and gas exchange system of the C57BL6 mice^24^ requires higher concentrations (85% O_2_) and a longer duration (P2-14) of oxygen exposure to induce human BPD-like lung pathology^9,25^. Our model, while not an exact simulation of the human preterm infant, reproduces both structurally the hippocampal shrinkage and functionally the associated memory deficits^26^ seen in adolescents and young adults with BPD.

The hippocampus, a region of the brain that plays an important role in consolidating short memory into long-term memory^14–16^, is highly vulnerable to oxidative stress^17^. Oxygen exposure causes neuronal cell death in developing brain^11,12^, and prolonged oxidative stress impairs neuronal mitochondrial function^13^. Neurons in the hippocampus are critically dependent on their mitochondrial function for the strengthening of synapses, a cellular response responsible for the formation and maintenance of long-term memory^27–29^. In neurons, mitochondria generate about 90% of the ATP by oxidative phosphorylation. In oxidative phosphorylation, oxygen is the terminal electron acceptor of the mitochondrial electron transport chain (ETC), which transfers electrons from high energy metabolites through a series of carriers to drive ATP generation from ADP^30^. The redox state of the respiratory chain is governed by the trans-membrane proton gradient and the membrane potential^31^. The redox energy used for ATP generation also leads to the production of reactive oxygen species (ROS)^32^. Excessive ROS production following hyperoxia exposure can potentially overwhelm antioxidant defense mechanisms and leads to mitochondrial damage^33,34^ and cellular death^35^.

Our mitochondrial functional assessments show that early life hyperoxia exposure not only reduces ATP linked oxygen consumption in the hippocampus in the neonatal period (P14) but also in the young adult (14 weeks). In addition, our study also shows a persistent increase in the rate of oxygen consumption at state 4 respiration, (a surrogate measure of proton leak) in young adults exposed to hyperoxia as neonates, and suggest uncoupling between substrate oxidation and ATP synthesis. Alterations in mitochondrial coupling can result in increased ROS production^36–38^ and impairment in ATP synthesis^39^. Though the amount of ATP produced by the hippocampal tissue was not measured in this study, the decrease in ATP linked oxygen consumption and increase in state 4 proton leak both at P14 and 14 weeks suggest that early life oxygen exposure permanently impairs mitochondrial efficiency in the generation of ATP. The neonatal hyperoxia-induced hippocampal mitochondrial dysfunction measured through bioenergetic studies in young adult mice is consistent with the mitochondrial dysfunction predicted through proteomic analysis.

Complex I (NADH: ubiquinone oxidoreductase), the first and largest enzyme in the ETC, has been consistently shown to be vulnerable to oxidative stress-mediated dysfunction^40^. It is also thought to be the main site of ROS production^41,42^, and its impairment leads to an increase in ROS production^43^. Decreased complex I activity seen in hyperoxia-exposed neonatal mice suggests that oxygen exposure either directly or indirectly impairs complex I function. At 14 weeks, the targeted hippocampal proteomic analysis determined decreases in complex I NDUFB8 and NDUFB11 subunits in neonatal hyperoxia-exposed mice. While neither of these subunits are thought to be directly involved in catalysis, reports have associated decreased levels of NDUFB8 in a mouse model of AD^44^. However, mitochondrial bioenergetic studies indicated an increase in complex I activity at 14 weeks. The persistent decreased ATP linked O_2_ consumption and increased state 4 proton leak at 14 weeks despite the increase in complex I activity in the young adult mice exposed to hyperoxia suggest persistent mitochondrial dysfunction and inadequate compensation by the later increase in complex I activity following hyperoxia-induced decreases in the newborn. Neonatal hyperoxia did not affect complex IV activity either at P14 or 14 weeks suggesting that either the complex IV enzyme is not as highly vulnerable to oxidative stress as complex I or it is well adapted to oxidative stress-induced injury. Though changes in hyperoxia-induced complex III and V activity are possible, comparable succinate-induced oxygen consumption between hyperoxia and air-exposed neonatal and young adult mice indicate that dysfunction in oxygen consumption noted with hyperoxia exposure are mainly induced by alterations in complex I function.

Mitochondrial metabolism and cell death signaling are sexually dimorphic. Compared to the female, the male hippocampus has a lower level of endogenous antioxidant defense systems^45^ and produces more ROS^46^. We determined that hyperoxia-exposed young males had reduced ATP linked O_2_ consumption and increased complex I activity compared to hyperoxia-exposed young females. These observations are clinically important because prematurity associated neurodevelopmental outcomes^47^ and neurodevelopmental disorders (e.g., Autism)^48^ preferentially affects male sex. Citrate synthase, a marker for mitochondrial volume^49^, is reduced by hyperoxia in neonates (P14) and normalized in young adults (14 weeks). This suggests that although the adult hippocampus is able to regenerate the early hyperoxia-induced mitochondrial loss, the regenerated mitochondria are still inefficient, as evident by the persistent mitochondrial dysfunction in the young adult mice.

In addition, hyperoxia-induced changes in the expression of Ras-related protein Rab-8A (involved in vesicular trafficking and neurotransmitter), Teneurin-1 (increases hippocampal dendritic arborization and spine density) and their roles in hyperoxia-induced cognitive dysfunction need further investigation. Our data also indicate that aberrant GABAergic signaling^50^ and amyloid processing are associated with cognitive deficits, and these pathways have been linked to neurodegenerative conditions^51^. Additional studies are needed to evaluate the contribution of these canonical pathways to impaired memory and hippocampal dysfunction induced by oxidative stress and to define how they interact with mitochondrial dysfunction.

## Conclusion

We have demonstrated that supraphysiological oxygen exposure during a critical developmental period has a permanent negative impact on hippocampal mitochondria. The pathophysiology of neonatal hyperoxia-induced permanent mitochondrial dysfunction is complex. We speculate that early oxidative stress increases mitochondrial ROS production through complex I dysfunction which in turn causes permanent damage to mitochondrial DNA leading to long-term alterations in complex I function and overall mitochondrial function (Supplemental Fig 2).

Future studies designed to quantitate mitochondrial DNA damage, ATP and ROS levels are needed to determine the mechanisms by which early hippocampal complex I dysfunction induces permanent complex I dysfunction and the development of spatial memory deficits.

## Materials and Methods

All protocols were approved by the UAB Institutional Animal Care and Use Committee (IACUC) and were consistent with the PHS Policy on Humane Care and Use of Laboratory Animals (Office of Laboratory Animal Welfare, Aug 2002) and the Guide for the Care and Use of Laboratory Animals (National Research Council, National Academy Press, 1996).

## Animal model

C57BL/6J dams and their pups of both sexes were exposed to either normobaric hyperoxia (85% O_2_, N=6) or normobaric 21% O_2_ ambient air (Air, N=6) from the second postnatal day (P2) until postnatal day 14 (P14), returned to room air, and maintained on standard rodent diet and light/dark cycling in microisolator cages until 14 weeks of age (Fig 4A)^25^. An additional set of mice were exposed to either 85% O_2_ (Hyperoxia, N=6) or 21% O_2_ (Air, N=6) and sacrificed at P14 (Fig 4A).

**Fig 4:**
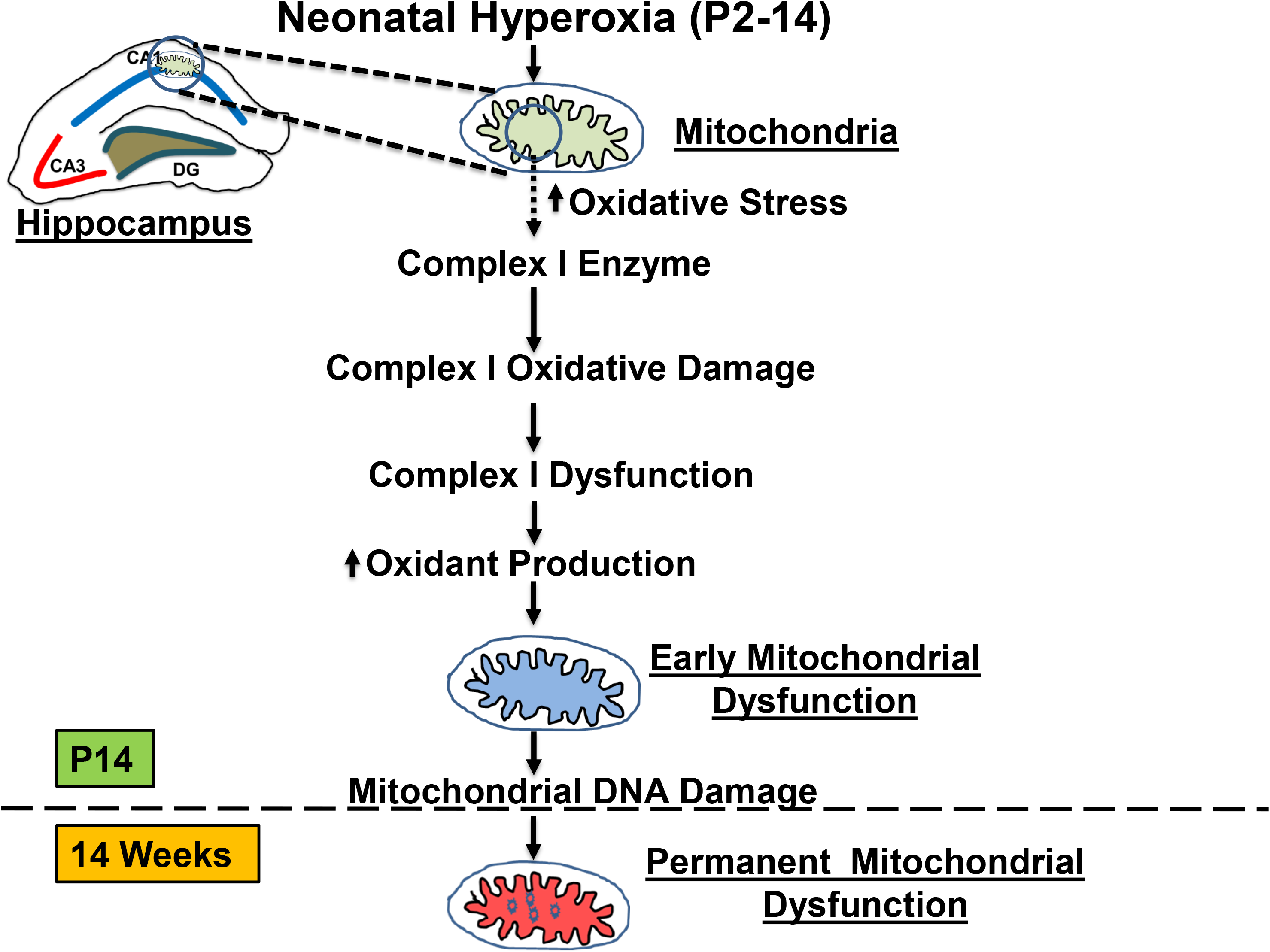
Schematics of the animal model and mitochondrial respiratory protocol. Fig A: Represents schematic of hyperoxia exposure from P2-14 and experimental studies done at P14 and 14 weeks. Fig B: Represents schematic of the mitochondrial respiratory protocol used in the high-resolution respirometry with sequentially added substrates and the calculations to assess the mitochondrial bioenergetic function using whole hippocampal tissue. ST2PM = State 2 Respiration with Pyruvate and Malate, ST3PM = State 3 Respiration with Pyruvate and Malate, ADP = Adenosine Diphosphate, ST3PMS = State 3 Respiration with Pyruvate, Malate and Succinate, ST4 = State 4 Respiration following Oligomycin, ATP Linked O_2_ = Adenosine triphosphate linked O_2_ consumption, ST4 Proton Leak= State 4 respiration proton leak, ROX = Residual O_2_ Consumption, and Baseline = Baseline O_2_ Consumption.

At 14 weeks, hippocampal proteins were analyzed by unbiased proteomic profiling using mass spectrometry. Initially, the targeted analysis was performed for hippocampal mitochondrial respiratory complex proteins. Subsequently, bioinformatics analysis was performed on all other differentially expressed hippocampal proteins between hyperoxia and air-exposed groups. At P14 and 14 weeks of age, hippocampal tissues were analyzed for mitochondrial bioenergetic functions, complex I, IV, and citrate synthase activities.

## Mass Spectrometry

At 14 weeks, following cervical dislocation, the whole brain was harvested, and hippocampi were removed in a sterile manner^10^. Tissue was then homogenized using Qiagen tissue lyser (Qiagen, MD, USA) in T-PER+Halt protease inhibitors+PMSF solution, and protein assay was performed using BCA protein assay kit (Thermo Fisher Scientific, MA, USA)^52^. The mass spectrometric analysis of hippocampal proteins was done as previously described^10^.

## Proteomics Data assessment

Differentially expressed proteins (fold change ±1.5 fold and p <0.05) were identified using T-test and further analyzed. As previously done^10^, functional analysis was performed using PANTHER (Protein ANalysis THrough Evolutionary Relationships)^53^ and Ingenuity Pathway Analysis (QIAGEN Inc. MD, USA). Heat maps were generated using pheatmap package V.1.0.7 in R program.

## Mitochondrial Bioenergetic Studies (High-resolution Respiratory)

Whole hippocampus (right) was harvested and placed in ice cold artificial cerebrospinal fluid that contained glucose, BSA, EGTA, pyruvate, and mitochondrial respiration buffer, as previously described^54^. Briefly, hippocampal tissue was permeabilized with saponin (5 mg/mL, 30 minutes) and high-resolution respirometry performed using a two-channel respirometer (Oroboros Oxygraph-2k with DatLab software; Oroboros, Innsbruck, Austria). Reactions were conducted at 37°C in a 2 ml chamber containing air-saturated mitochondrial respiration buffer (MiR03) under continuous stirring.

As illustrated in Fig 4B, O_2_ consumption rates were measured in the presence of substrates (5 mM malate, 15 mM pyruvate, 2.5mM ADP,10 mM succinate), and inhibitors (0.5 μM oligomycin, 5um antimycin A) to assess state 2 (substrate alone), state 3 (substrate + ADP) and oligomycin induced state 4 respiration rates. Non-mitochondrial oxygen consumption was determined in the presence of antimycin A. Adenosine triphosphate (ATP) linked O_2_ consumption rate was determined by State 3 (substrates + ADP) - State 4 (oligomycin) = ATP linked rate. Non-ATP linked O_2_ consumption rate was determined by State 4 (oligomycin) – non-mitochondrial oxygen consumption (antimycin A). Potential differences in O_2_ consumption rates based upon substrate utilization at complex I or II were assessed using pyruvate/malate or succinate, respectively, in the presence of ADP.

## Complex I, IV and Citrate Synthase Activity Assays

Complex I, IV, and citrate synthase activities were measured from hippocampal (left) homogenates as previously described^55–57^; complex I activities were measured from freshly extracted tissues.

### Statistical Analysis

Results were expressed as means ±SE. Multiple comparisons testing (Student-Newman-Keuls) was performed if statistical significance (*p* < 0.05) was noted by ANOVA.

## Acknowledgments

The authors wish to thank the UAB BioAnalytical Redox Biology Core (P30DK079626) and Dr. Douglas Moellering for assistance in the mitochondrial functional measures. This work was partially funded by R01HL092906, R01AG021612, R01NS076312, R25NS089463, and the Kaul Pediatric Research Institute, Division of Neonatology, Department of Pediatrics, University of Alabama at Birmingham, Alabama USA.

## Author contributions

Concept and design: MR, LLM, SWB, and NA; Data analysis and interpretation: MR, RK JK, LLM, SWB and NA; Drafting the manuscript for important intellectual content: MR, SWB, and NA. Performed the research: MR, and KM. All authors edited and approved the final version of the manuscript.

---

**Supplemental Fig 1:**

Hierarchical clustering of differentially expressed hippocampal proteins in room air vs hyperoxia exposed young adult mice. Dendrogram above the heat map depicts hierarchical clustering of the samples (Room air is shown as red, hyperoxia samples cyan). Cluster distance is based on the average distance between all the pairs of objects in the two clusters. Dendrograms for differentially expressed proteins are shown on the left side of the heat map. In the heat maps, red shows increased expression, and blue indicates decreased expression. Expression value intensities are illustrated by color with a range of −2 to 8 on a log scale.

**Supplemental Fig 2:**

Cartoon depicting the possible mechanism(s) by which early life oxygen exposure leads to permanent hippocampal mitochondrial dysfunction.

