## Supplemental Table 1 for "Early Life Supraphysiological Levels of Oxygen Exposure Permanently Impairs Hippocampal Mitochondrial Function"

**Abbreviated title:** Oxygen-Induced Hippocampal Mitochondrial Dysfunction

**Author names and affiliations:**

*Manimaran Ramani MD^1^, Kiara Miller, BS^1^, Ranjit Kumar PhD^2^, Jegen Kadasamy^1^, Lori McMahon PhD^3,4^, Scott Ballinger PhD^5^_,_ and Namasivayam Ambalavanan MD^1, 2^

From the Departments of Pediatrics^1^, Bioinformatics^2^, Cell, Developmental, and Integrative Biology^3^, Neurobiology^4^, and Pathology^5^ University of Alabama at Birmingham, Birmingham, AL 35233

**Corresponding author:**

*Manimaran Ramani MD, University of Alabama at Birmingham, 176F Suite 9380,

**Supplemental Table 1**: Upregulated Hippocampal Proteins in Young Adult Mice Exposed to Neonatal Hyperoxia (n=5 in Air group, 5 in Hyperoxia group)

| **Molecule (Symbol)** | **Protein**  **Log Fold Change in Hyperoxia (vs. Air)** | **P value for protein change** |
| --- | --- | --- |
| Ras-related protein Rab-8A (RAB8A) | + 4.78 | 0.032 |
| Cytochrome b-c1 complex subunit 9 (UQCR9) | +4.44 | 0.0005 |
| Regulator complex protein LAMTOR3 (LTOR3) | +4.35 | 0.02 |
| 39S ribosomal protein L11, mitochondrial (RM11) | +4.19 | 0.003 |
| Myelin proteolipid (PLP) | +4.05 | 0.003 |
| Protein RRP5 homolog (RRP5) | +3.88 | 0.02 |
| Peptidyl-prolyl cis-trans isomerase F, mitochondrial (PPIF) | +3.87 | 0.003 |
| Otoferlin (OTOF) | +3.74 | 0.01 |
| Vesicle-associated membrane protein 7 (VAMP7) | +3.63 | 0.02 |
| Guanine nucleotide-binding protein subunit beta-4 (GBB4) | +3.61 | 0.04 |
| 28S ribosomal protein S21, mitochondrial (RT21) | +3.60 | 0.01 |
| Transducin-like enhancer protein 3 (TLE3) | +3.54 | 0.02 |
| Translocon-associated protein subunit delta (SSRD) | +3.51 | 0.004 |
| Kinesin-associated protein 3 (KIFA3) | +3.47 | 0.01 |
| Glutamate receptor ionotropic, kainate 3 (GRIK3) | +3.47 | 0.01 |
| SRSF protein kinase 2 (SRPK2) | +3.46 | 0.01 |
| Lysine-specific histone demethylase 1A (KDM1A) | +3.44 | 0.03 |
| GES30 (Q9Z2P7) | +3.42 | 0.01 |
| Low-density lipoprotein receptor-related protein 4 (LRP4) | +3.42 | 0.02 |
| Nuclear valosin-containing protein-like (NVL) | +3.33 | 0.03 |
| Shugoshin-like 2 (SGOL2) | +3.33 | 0.04 |
| Target of Myb protein 1 (TOM1) | +3.20 | 0.01 |
| Casein kinase II subunit alpha (CSNK2A1) | +3.14 | 0.02 |
| Dynein light chain Tctex-type 1 (DYLT1) | +2.98 | 0.02 |
| Inositol-trisphosphate 3-kinase A (IP3KA) | +2.93 | 0.03 |
| Peroxisomal biogenesis factor 19 (PEX19) | +2.92 | 0.01 |
| Galectin-related protein (LEGL) | +2.86 | 0.03 |
| CDP-diacylglycerol--inositol 3-phosphatidyltransferase (CDIPT) | +2.86 | 0.02 |
| E3 ubiquitin-protein ligase RNF19A (RN19A) | +2.84 | 0.02 |
| 60S ribosomal protein L22-like 1 (RL22L) | +2.82 | 0.02 |
| Eukaryotic translation initiation factor 3 subunit F (EIF3F) | +2.79 | 0.04 |
| von Willebrand factor A domain-containing protein 8 (VWA8) | +2.78 | 0.05 |
| Exosome complex exonuclease RRP44 (RRP44) | +2.70 | 0.03 |
| Thioredoxin, mitochondrial (THIOM) | +2.61 | 0.01 |
| Magnesium-dependent phosphatase 1 (MGDP1) | +2.53 | 0.03 |
| Retinaldehyde-binding protein 1 (RLBP1) | +2.53 | 0.03 |
| Armet protein (AMERT) | +2.53 | 0.05 |
| Chromatin target of PRMT1 protein (CHTOP) | +2.43 | 0.05 |
| Ubiquitin carboxyl-terminal hydrolase 11 (UBP11) | +2.43 | 0.01 |
| 28S ribosomal protein S28, mitochondrial (RT28) | +2.41 | 0.03 |
| Glutamate decarboxylase 1 (DCE1) | +2.24 | 0.04 |
| Cytochrome b-5, isoform CRA_a (CYB5B) | +2.10 | 0.05 |
| Fatty acid desaturase 2 (FADS2) | +2.04 | 0.05 |
| Cysteine desulfurase, mitochondrial (NFS1) | +1.98 | 0.0005 |
| Autophagy protein 5 (ATG5) | +1.96 | 0.03 |
| Titin (TITIN) | +1.93 | 0.03 |
| Protein-L-isoaspartate O-methyltransferase (PCMT1) | +1.51 | 0.03 |
| Interferon-induced very large GTPase 1 (GVIN1) | +1.50 | 0.04 |
