## Supplemental Table 2 for "Early Life Supraphysiological Levels of Oxygen Exposure Permanently Impairs Hippocampal Mitochondrial Function"

**Abbreviated title:** Oxygen-Induced Hippocampal Mitochondrial Dysfunction

**Author names and affiliations:**

*Manimaran Ramani MD^1^, Kiara Miller, BS^1^, Ranjit Kumar PhD^2^, Jegen Kadasamy^1^, Lori McMahon PhD^3,4^, Scott Ballinger PhD^5^_,_ and Namasivayam Ambalavanan MD^1, 2^

From the Departments of Pediatrics^1^, Bioinformatics^2^, Cell, Developmental, and Integrative Biology^3^, Neurobiology^4^, and Pathology^5^ University of Alabama at Birmingham, Birmingham, AL 35233

**Corresponding author:**

*Manimaran Ramani MD, University of Alabama at Birmingham, 176F Suite 9380,

**Supplemental Table 2:** Downregulated Hippocampal Proteins in Young Adult Mice Exposed to Neonatal Hyperoxia (n=5 in Air group, 5 in Hyperoxia group)

| **Molecule (Symbol)** | **Protein**  **Log Fold Change in Hyperoxia (vs. Air)** | **P value for protein change** |
| --- | --- | --- |
| Filamin-C (FLNC) | -5.13 | 9.97E-07 |
| Vacuolar Protein Sorting-Associated Protein 52 Homolog (VPS52) | -4.74 | 1.79E-06 |
| Integrin Beta-1 (ITB1) | -4.50 | 5.77E-07 |
| Glucose-6-Phosphate 1-Dehydrogenase X (G6PD1) | -4.44 | 0.01 |
| Teneurin-1 (TEN1) | -4.38 | 0.0001 |
| Serine/Threonine-Protein Kinase (MARK1) | -4.17 | 0.01 |
| CCA Trna Nucleotidyltransferase 1(TRNT1) | -4.11 | 9.86E-05 |
| Actin-Like Protein 6B (ACL6B) | -4.07 | 0.001 |
| Cold Shock Domain-Containing Protein E1 (CSDE1) | -4.05 | 0.05 |
| RNA Binding Protein Fox-1 Homolog 2 (RFOX2) | -4.01 | 0.01 |
| Coatomer Subunit Beta 2 (COPB2) | -4.01 | 0.02 |
| E3 Ubiquitin-Protein Ligase HECW1 (HECW1) | -3.97 | 2.16E-05 |
| Nuclear Mitotic Apparatus Protein 1 (NUMA1) | -3.88 | 0.01 |
| Melanoma-Associated Antigen D1 (MAGD1) | -3.87 | 4.92E-05 |
| Glutaminyl-Trna Synthetase, Isoform CRA_A (QARS) | -3.87 | 0.01 |
| U2 Snrnp-Associated SURP Motif-Containing Protein (SR140) | -3.86 | 0.01 |
| Y-Box-Binding Protein 3 (YBOX3) | -3.85 | 0.03 |
| Phosphatidate Cytidylyltransferase 2 (CDS2) | -3.83 | 0.02 |
| CDE1-Binding Protein (CDEBP) | -3.83 | 0.003 |
| AFG3-Like Protein 2 (AFG32) | -3.82 | 0.01 |
| Axin Interactor, Dorsalization-Associated Protein  (AIDA) | -3.79 | 0.01 |
| Ubiquitin Conjugation Factor E4 B (UBE4B) | -3.78 | 0.004 |
| WD Repeat-Containing Protein 6 (WDR6) | -3.78 | 4.21E-05 |
| Acetoacetyl-Coa Synthetase (AACS) | -3.76 | 0.003 |
| Nucleobindin-1 (NUCB1) | -3.76 | 0.01 |
| Endoplasmic Reticulum-Golgi Intermediate Compartment Protein 1 (ERGI1) | -3.75 | 0.01 |
| Fibronectin (FINC) | -3.74 | 0.01 |
| Eukaryotic Translation Initiation Factor 3 Subunit K (EIF3K) | -3.68 | 0.01 |
| ELAV-Like Protein 2 (ELAV2) | -3.68 | 0.04 |
| Striatin-4 (STRN4) | -3.64 | 0.02 |
| Myosin-14 (MYH14) | -3.63 | 0.03 |
| Acidic Leucine-Rich Nuclear Phosphoprotein 32 Family Member E (AN32E) | -3.63 | 0.03 |
| Synaptic Vesicle Membrane Protein VAT-1 Homolog-Like (VAT1L) | -3.62 | 0.04 |
| CTP Synthase 1 (PYRG1) | -3.61 | 0.001 |
| 39S Ribosomal Protein L15, Mitochondrial  (RM15) | -3.60 | 0.004 |
| Protein Tweety Homolog 3 (TTYH3) | -3.60 | 0.003 |
| Coiled-Coil Domain-Containing Protein 47 (CCD47) | -3.53 | 0.003 |
| Beta-Arrestin-1 (ARRB1) | -3.53 | 0.01 |
| Kinesin-Like Protein KIF21A (KI21A) | -3.50 | 0.05 |
| NACHT And WD Repeat Domain-Containing Protein 2 (NWD2) | -3.47 | 0.01 |
| Splicing Factor 3B Subunit 3 (SF3B3) | -3.47 | 0.02 |
| Neuron Navigator 3 (NAV3) | -3.46 | 0.04 |
| Golgin Subfamily A Member 4 (GOGA4) | -3.42 | 0.01 |
| RNA-Binding Protein 10 (RBM10) | -3.42 | 0.004 |
| Oligodendrocyte-Myelin Glycoprotein (OMGP) | -3.37 | 0.01 |
| Rab3 Gtpase-Activating Protein Non-Catalytic Subunit (RBGPR) | -3.36 | 0.03 |
| Golgin Subfamily A Member 2 (GOGA2) | -3.35 | 0.01 |
| Splicing Factor 3B Subunit 1 (SF3B1) | -3.32 | 0.02 |
| Ras Association and Pleckstrin Homology Domains 1 (RAPH1) | -3.31 | 0.01 |
| Brain-Specific Angiogenesis Inhibitor 3 (BAI3) | -3.30 | 0.02 |
| Protein ADP-Ribosylarginine Hydrolase (ADPRH) | -3.29 | 0.04 |
| rRNA 2'-O-Methyltransferase Fibrillarin (FBRL) | -3.28 | 3.61E-05 |
| Ubiquitin Conjugation Factor E4 A (UBE4A) | -3.26 | 0.01 |
| Protein Transport Protein Sec31B (SC31B) | -3.25 | 0.01 |
| Striatin-Interacting Protein 1 (STRP1) | -3.25 | 0.004 |
| Voltage-Dependent R-Type Calcium Channel Subunit Alpha-1E (CAC1E) | -3.24 | 0.03 |
| Apolipoprotein O (APOO) | -3.22 | 0.01 |
| Serine/Threonine-Protein Kinase MARK2 (MARK2) | -3.20 | 0.02 |
| Dedicator Of Cytokinesis Protein 11 (DOC110 | -3.19 | 0.01 |
| Gtpase-Activating Protein And VPS9 Domain-Containing Protein 1 (GAPD1) | -3.17 | 0.03 |
| Regulating Synaptic Membrane Exocytosis Protein 1 (RIMS1) | -3.17 | 0.04 |
| Clathrin Interactor 1 (CLINT1) | -3.16 | 0.02 |
| Hepatocyte Growth Factor-Regulated Tyrosine Kinase Substrate (HGS) | -3.16 | 0.03 |
| Asparagine Synthetase (ASNS) | -3.13 | 0.02 |
| La-Related Protein 6 (LARP6) | -3.12 | 0.01 |
| Elongation Factor Tu GTP Binding Domain Containing 2 (EFTUD2) | -3.11 | 0.04 |
| N-Acetyl-D-Glucosamine Kinase (NAGK) | -3.11 | 0.04 |
| Integrin Alpha-V (ITAV) | -3.07 | 0.03 |
| Syntaxin-6 (STX6) | -3.07 | 0.03 |
| Huntingtin-Interacting Protein 1(HIP1) | -3.05 | 0.04 |
| O-Acetyl-ADP-Ribose Deacetylase MACROD2 (MACD2) | -3.05 | 0.01 |
| Neuropilin-1 (NRP1) | -3.04 | 0.04 |
| Integrator Complex Subunit 1 (INT1) | -3.03 | 0.03 |
| Protein FAM171A2 (F1712) | -3.01 | 0.02 |
| Small G Protein Signaling Modulator 1 (SGSM1) | -2.99 | 0.02 |
| Probable ATP-Dependent RNA Helicase DDX10  (DDX10) | -2.97 | 0.01 |
| Ankyrin Repeat Domain-Containing Protein 17 (ANR17) | -2.97 | 0.02 |
| Proteasome Activator Complex Subunit 4  (PSME4) | -2.95 | 0.03 |
| Stathmin-4 (STMN4) | -2.94 | 0.05 |
| Syntenin-1 (SDCB1) | -2.92 | 0.04 |
| Methylcrotonoyl-Coa Carboxylase Subunit Alpha, Mitochondrial (MCCA) | -2.92 | 0.02 |
| Voltage-Dependent Calcium Channel Subunit Alpha-2/Delta-3 (CA2D3) | -2.92 | 0.03 |
| Plakophilin-4 (PKP4) | -2.91 | 0.03 |
| Gamma-Aminobutyric Acid Receptor Subunit Alpha-5 (GBRA5) | -2.90 | 0.03 |
| Copper Chaperone For Superoxide Dismutase (CCS) | -2.90 | 0.02 |
| Actin-Related Protein 10 (ARP10) | -2.89 | 0.03 |
| TBC1 Domain Family Member 24 (TBC24) | -2.89 | 0.03 |
| OL-Protocadherin Isoform (PCDH10) | -2.89 | 0.05 |
| SAPS Domain Family, Member 3, Isoform CRA_C (SAPS3) | -2.88 | 0.05 |
| Putative GTP Cyclohydrolase 1 Type 2 Nif3l1 (NIF3L1) | -2.88 | 0.04 |
| Proline-Rich and Coiled-Coil-Containing Protein 2C (PRC2C) | -2.87 | 0.05 |
| ADP-Ribose Pyrophosphatase, Mitochondrial (NUDT9) | -2.86 | 0.03 |
| Annexin A2 (ANXA2) | -2.86 | 0.04 |
| Phosphofurin Acidic Cluster Sorting Protein 2 (PACS2) | -2.85 | 0.03 |
| Probable Trna N6-Adenosine Threonylcarbamoyltransferase (OSGEP) | -2.82 | 0.03 |
| Gamma-Aminobutyric Acid Receptor Subunit Alpha-3  (GBRA3) | -2.80 | 0.04 |
| YTH Domain-Containing Family Protein 3 (YTHD3) | -2.80 | 0.02 |
| Splicing Factor 45 (SPF45) | -2.80 | 0.03 |
| Ubiquitin Carboxyl-Terminal Hydrolase 24 (UBP24) | -2.79 | 0.03 |
| Host Cell Factor 1 (HCFC1) | -2.79 | 0.05 |
| Adenylate Cyclase Type 5 (ADCY5) | -2.78 | 0.03 |
| Immunoglobulin Superfamily Member 3 (IGSF3) | -2.77 | 0.03 |
| 28S Ribosomal Protein S6, Mitochondrial (RT06) | -2.76 | 0.03 |
| Zinc Finger Protein 638 (ZN638) | -2.76 | 0.01 |
| Rab Gtpase-Activating Protein 1 (RBGP1) | -2.76 | 0.03 |
| SWI/SNF-Related Matrix-Associated Actin-Dependent Regulator of Chromatin Subfamily D Member 3 (SMRD3) | -2.75 | 0.01 |
| La-Related Protein 1 (LARP1) | -2.75 | 0.03 |
| Ran-Binding Protein 3 (RANB3) | -2.72 | 0.03 |
| Activity-Dependent Neuroprotector Homeobox Protein (ADNP) | -2.71 | 0.004 |
| Phosphatidylinositol 3,4,5-Trisphosphate 3-Phosphatase And Dual-Specificity Protein Phosphatase (PTEN) | -2.71 | 0.05 |
| Alpha/Beta Hydrolase Domain-Containing Protein 11 (ABHDB) | -2.70 | 0.02 |
| Epidermal Growth Factor Receptor Kinase Substrate 8 (EPS8) | -2.70 | 0.03 |
| Myotubularin-Related Protein 9 (MTMR9) | -2.69 | 0.02 |
| Deoxyhypusine Hydroxylase (DOHH) | -2.63 | 0.03 |
| Inter-Alpha-Trypsin Inhibitor Heavy Chain H2 (ITIH2) | -2.60 | 0.03 |
| Serine/Threonine-Protein Kinase TAO2 (TAOK2) | -2.60 | 0.03 |
| Integrator Complex Subunit 3 (INT3) | -2.56 | 0.03 |
| YLP Motif-Containing Protein 1 (YLPM1) | -2.55 | 0.03 |
| Glucosamine-6-Phosphate Isomerase 2 (GNPI2) | -2.54 | 0.03 |
| Casein Kinase I Isoform Alpha (H7BXB1) | -2.53 | 0.03 |
| Dystroglycan (DAG1) | -2.53 | 0.03 |
| Protein Transport Protein Sec24A (SC24A) | -2.53 | 0.03 |
| Syndecan-3 (SDC3) | -2.53 | 0.03 |
| Serine/Threonine-Protein Kinase LMTK3 (LMTK3) | -2.48 | 0.03 |
| Potassium Voltage-Gated Channel Subfamily D Member 2 (KCND2) | -2.47 | 0.03 |
| NAD(P) Transhydrogenase, Mitochondrial (NNTM) | -2.43 | 0.03 |
| Calcium-Transporting Atpase Type 2C Member 1 (AT2C1) | -2.43 | 0.03 |
| Nuclear Autoantigenic Sperm Protein (NASP) | -2.41 | 0.03 |
| NADH Dehydrogenase [Ubiquinone] 1 Beta Subcomplex Subunit 11, Mitochondrial (NDUBB) | -2.37 | 0.03 |
| SRSF Protein Kinase 1 (SRPK1) | -2.37 | 0.03 |
| Quinone Oxidoreductase-Like Protein 1 (QORL1) | -2.34 | 0.03 |
| Golgin Subfamily A Member 3 (GOGA3) | -2.31 | 0.05 |
| Roundabout Homolog 1 (ROBO1) | -2.30 | 0.04 |
| Protein Daple (Daple) | -2.26 | 0.03 |
| Betacstf-64 Variant 3 (B3V097) | -2.25 | 0.04 |
| Annexin A1 (ANXA1) | -2.25 | 0.03 |
| Hydrocephalus-Inducing Protein (HYDIN) | -2.24 | 0.04 |
| Alpha-Methylacyl-Coa Racemase (AMACR) | -2.15 | 0.03 |
| Serine/Threonine-Protein Phosphatase CPPED1 (CPPED) | -2.13 | 0.03 |
| Girdin (GRDN) | -1.76 | 0.02 |
| Acetyl-Coa Carboxylase 1 (ACACA) | -1.75 | 0.04 |
| Chondroitin Sulfate Proteoglycan 4 (CSPG4) | -1.56 | 0.01 |
